# Consistency, Inconsistency and Ambiguity of Metabolite Names in Biochemical Databases Used for Genome Scale Metabolic Modelling

**DOI:** 10.1101/503664

**Authors:** Nhung Pham, Ruben G. A. van Heck, Jesse C. J. van Dam, Peter J. Schaap, Edoardo Saccenti, Maria Suarez-Diez

## Abstract

Genome scale metabolic models (GEMs) are manually curated repositories describing the metabolic capabilities of an organism. GEMs have been successfully used in different research areas, ranging from systems medicine to biotechnology. However, the different naming conventions (namespaces) of databases used to build GEMs limit model reusability and prevent the integration of existing models. This problem is known in the GEM community but its extent has not been analyzed in depth. In this study, we investigate the name ambiguity and the multiplicity of non-systematic identifiers and we highlight the (in)consistency in their use in eleven biochemical databases of biochemical reactions and the problems that arise when mapping between different namespaces and databases. We found that such inconsistencies can be as high as 83.1%, thus emphasizing the need for strategies to deal with these issues. Currently, manual verification of the mappings appears to be the only solution to remove inconsistencies when combining models. Finally, we discuss several possible approaches to facilitate (future) unambiguous mapping.

## 1. Introduction

Genome scale metabolic models (GEMs) combine available metabolic knowledge of an organism in a consistent and structured way that allows prediction and simulation of metabolic phenotypes [1]. GEMs have been successfully used in different research areas, ranging from biotechnology to systems medicine, often resulting in new insights on metabolic processes in living organisms [2–5]. GEMs may differ in content and scope, and can contain anything from a few hundred to a few thousand reactions and metabolites. However, the structure of the model remains similar regardless of the application: the main components are metabolites, metabolic reactions, enzymes and the corresponding encoding genes.

The construction of a GEM includes three main steps [6,7], First, the genome of the organism considered is functionally annotated in order to identify enzymes and the associated reactions and metabolites. Second, the list of enzymes and reactions is converted into a mathematical model, a so-called draft model, in the form of a stoichiometric matrix to which constraints are added to account for reaction reversibility and uptake and secretion of metabolites. Last, the model is manually curated using experimental data (such as growth data), information from literature and/ or expert knowledge. Manual curation involves human workload and entails the verification of each reaction in the model and its corresponding constraints, which is a very time-consuming task. Tools and pipelines (such as, for example, the SEED [8], Pathway Tools [9], and the Raven toolbox [10]) have been developed to automatize the annotation, draft the reconstruction and to aid high-throughput creation of genome scale draft models [11].

The tools for automated draft reconstruction rely on biochemical databases that are used to find reactions associated to the enzymes identified in the genome through annotation. In general, different tools use different databases. For instance: the SEED uses its own naming system [8], Pathway Tools [9] uses MetaCyc [12], and Raven [10] uses KEGG [13]. Every database uses its own namespace which is a particular set of identifiers (such as numerical tags or names) for metabolites and reactions: because of this, it can happen that the same metabolites and reactions have different naming conventions when different tools are used to generate draft GEMs. To complicate the matter further, researchers often tend to use their own naming conventions such as custom abbreviations for metabolites or consecutive numbering for metabolites and reactions and this adds up to the observed heterogeneity of names and identifiers found in GEMs available in the literature [14]: the use of unique identifiers, independent from the particular databases used, such as InchI [15,16], or references to interlink different namespaces, have been suggested as an essential and fundamental part of GEM [17] but this is seldom implemented.

GEMs are manually curated knowledge repositories integrating information from independent (organism-specific) sources and thereby provide a comprehensive representation of what is presently known about the metabolism of the modelled organism. There is often the need to combine the information stored in individual GEMs to arrive to a consensus metabolic model for a given organism [18,19]. The use of different namespaces limits the reusability of a GEM and often makes it impossible, or extremely laborious, to combine two GEMs. Further, it often hampers model expansion, which is the addition of new reactions and/or metabolites to an existing model because if different namespaces are used the same metabolite can be added many times with different names and, as a consequence, considered as different chemical entities which can, in the worst case, invalidate the model.

In principle, different GEMs can be combined into a community model (partially) representing the different organisms present in a microbial community, with the aim of modelling community metabolic interactions such as cross feeding or substrate competition [20]. Since mapping manually different namespaces is highly laborious and practically unfeasible for large models [18], the only viable solution to integrate different GEMs has often been to rebuild the required models *de novo* [21,22]. However, while this approach leads to models that can be easily combined, it causes the loss of all the expert knowledge introduced in the manual curation process.

Naive direct comparison of names using string algorithms is often insufficient [23] and to help mapping among different namespaces in a more systematic way tools for consensus model generation and for automatic translation have been introduced [19,24], together with databases like MNXRef from MetaNetX [25] and MetRxn [26], developed to provide cross-linking among the identifiers in the namespaces of different databases.

As a matter of fact, mapping different namespaces using metabolite or reaction identifiers is not a trivial task because researchers often refer to compounds with many different names and abbreviations and the namespaces reflect this (Figure 1A). Often in GEMs different chemical entities (like, for instance, citrate and citric acid) are used as exchangeable names and may end up in databases like BIGG (which harvest reactions which have been used in metabolic modelling) resulting in imprecise, misleading and sometimes incorrect synonyms. Similarly, GEMs are often built featuring reactions using generic compound classes (such as ‘Lipids’ or ‘Protein’). When these are included in GEMs databases they cause the same compound to be linked to different identifiers.

**Figure 1.**
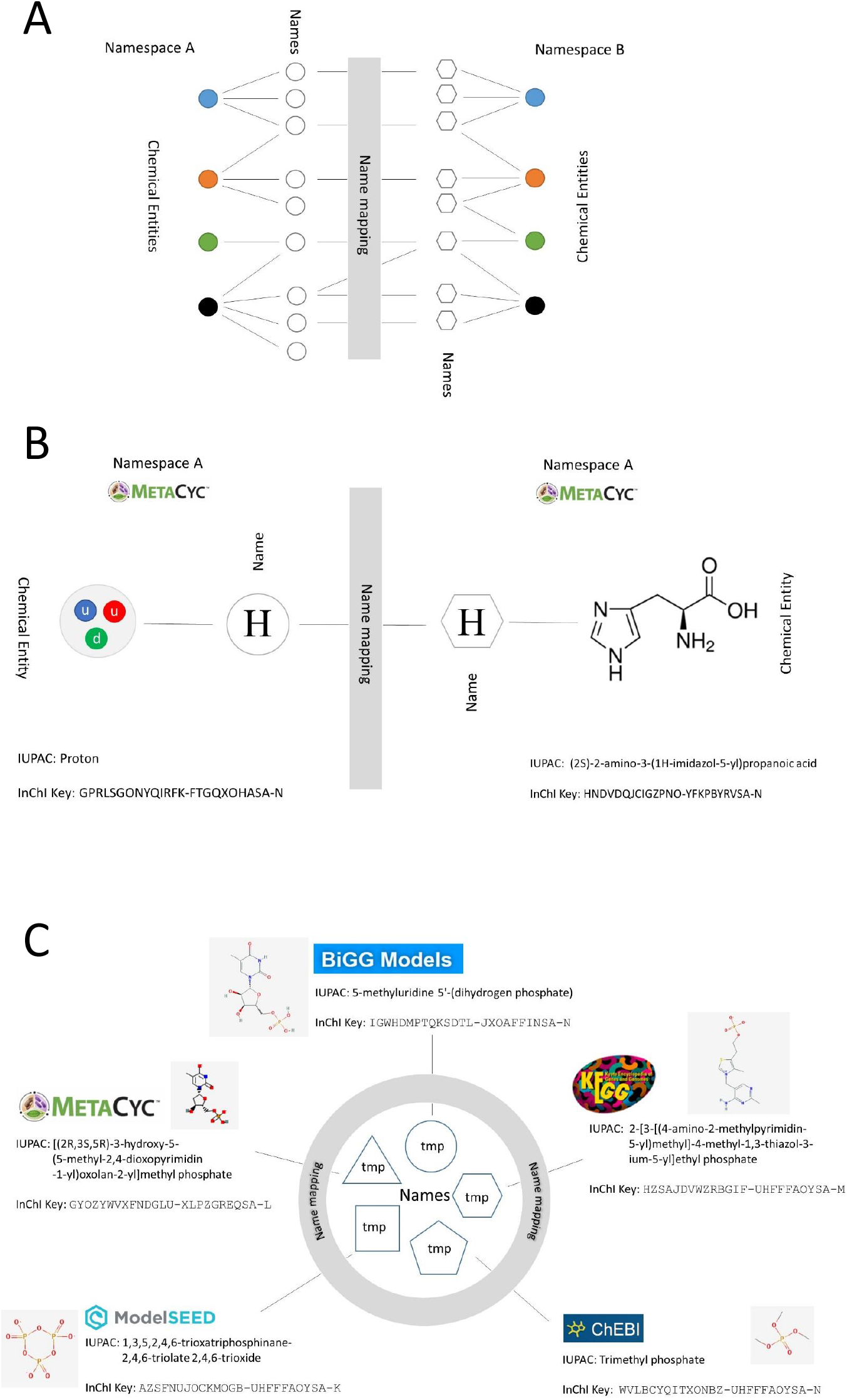
Inconsistent namespace problem statements. **A.** The same chemical entities (colored nodes) link to different names (colorless nodes) in different namespaces. Names in namespace A may link to different chemical entities in namespace B; **B.** Example of inconsistency within the same namespace, the same name links to different chemical compounds; **C.** Example of inconsistency between different namespaces. The same name links to different compounds in different databases. Chemical entities are represented with colored nodes, names are represented with colorless nodes

Internal database inconsistency is also often caused by ambiguous abbreviations, with the same shorthand used for different compound (Figure 1B). To make the matter worse, the same abbreviation can refer to different compounds in different databases (see Figure 1C).

The problems deriving from the inconsistency and the ambiguity in the namespaces of reaction databases used to build GEMs have been mentioned before [27–30] and are a well-known source of complaints in the modelling community. However, since the extent of the namespace mapping problem has not been so far analyzed in depth, we investigate the level of inconsistency and ambiguity encountered when i) mapping metabolites within a database and *ii*) mapping metabolites between two databases. To this task, we analyzed and compared naming and identifier conventions in eleven biochemical databases commonly used for metabolic modelling and metabolomics data analysis. Similar research has been done for small-molecule databases but did not considered databases used for metabolic modelling [31]. With this work we aim at raising awareness on this problem within the modelling community; provide a framework for evaluating when (or whether) GEMs and databases can be combined, suggest practices for dealing with this issue on the short term and, outline a strategy for a long term solution.

## 2. Results

To avoid ambiguity, we explicitly define the more specific terms used in this study as follows:

- *Identifier (ID)*: Identifiers are strings of alpha-numeric characters used to identify uniquely a metabolite or a reaction in a database. Examples are C00001 in KEGG or ATPM in BIGG.
- *Name*: Here we use name to refer not only to the chemical name, but also to the set of aliases, synonyms and abbreviations that are often included in a database as other names of the compound. For instance the KEGG ID C00001 is associated to the name ‘water’.
- *Multiplicity*: describes the case on which a single ID is linked to multiple names. For instance, The KEGG ID C00001 is associated to the names ‘water’ and ‘H20’; therefore we state that this ID has a multiplicity of 2.
- *Ambiguous*. The Merriam-Webster dictionary defines ambiguous (second entry) as ‘capable of being understood in two or more possible senses or ways’. Here, we use ambiguous (and its derivatives) to refer to the case on which the same name links to more than one ID in the same database. An example is shown in Figure 1 B, where the name ‘H’ links to the MetaCyc IDs ‘PROTON’ and ‘HIS’, associated to ‘hydrogen ion’ and ‘L-histidine’, respectively.
- *Consistency:* We use consistency (and its derivatives like consistent) to refer to mappings on which a molecular entity is mapped to itself. It follows that inconsistency is used to indicate a mapping or a database on which a molecular entity is associated to a different one.

We have analyzed eleven biochemical databases for their consistency and we have performed pairwise comparisons to investigate the degree of inter-database consistency. These databases were chosen for this study, primarily, because they were integrated in MetaNetX which facilitates data retrieval. In addition, many of them, namely BiGG, KEGG, SEED, HMDB, ChEBI and MetaCyc are databases often used for metabolic model reconstruction [14,32].

### 2.1. Mappings within the same database

#### 2.1.1. Name ambiguity

We calculated the average number of IDs per compound name for each of the eleven databases: results are summarized in Table 1.

**Table 1.** Ambiguity in biochemical database: number of names associated to more than one ID.

| Database | #Name | Average number of IDs per name<br>± standard deviation | % Ambiguous<br>names | # Ambiguous<br>names | Highest number<br>of IDs per name |
| --- | --- | --- | --- | --- | --- |
| BiGG | 5102 | 1.0 ± 0.1 | 1.31 | 67 | 3 |
| ChEBI | 388505 | 1.4 ± 1.5 | 14.8 | 57497 | 413 |
| enviPath | 11648 | 1.1 ± 0.3 | 7.38 | 860 | 10 |
| HMDB | 101101 | 1.0 ± 3.9 | 1.67 | 1686 | 921 |
| KEGG | 59682 | 1.1 ± 0.4 | 13.3 | 7936 | 16 |
| LIPID MAPS | 77457 | 1.0 ± 0.3 | 0.62 | 478 | 63 |
| MetaCyc | 55823 | 1.0 ± 0.1 | 0.5 | 279 | 13 |
| Reactome | 6972 | 1.8 ± 2.5 | 29.43 | 2052 | 34 |
| SABIO-RK | 11475 | 1.0 ± 0.0 | 0.07 | 8 | 3 |
| SEED | 47410 | 1.0 ± 0.1 | 1.06 | 503 | 4 |
| SLM | 1218750 | 1.1 ± 0.3 | 6.72 | 81894 | 9 |

With ChEBI and Reactome as exceptions, in most databases the average ID number is around 1: however there is a low consistency. Reactome has the lowest consistency: nearly 30% of compounds are associated with more than one ID, metabolites with generic descriptive names like ‘secretory granule lumen proteins’, ‘secretory granule membrane proteins’, and ‘ficolin-rich granule lumen proteins’ associate to 34 different IDs; there are also more specific names, like ‘hydron’, ‘water’ and ‘ATP’ associated to 21, 14 and 11 IDs, respectively. In the latter cases the cause is that different IDs are used to indicate the same metabolite in different subcellular compartments, although they all get assigned to the same name, for example the ID 5278291 indicates water in the cytoplasm while water in extracellular compartment is identified as 109276.

Overall, the most ambiguous metabolite name is ‘lecithin’, which is associated to 921 different IDs in the Human Metabolome database (HMDB). In this database, the most ambiguous names are general compound classes such as ‘diacylglycerol’, ‘PPP’ and ‘pyridin-3-ylboronic acid’.

The overall consistency of HMDB is very high, as only 1.7% of names are linked to multiple IDs, followed by ChEBI and KEGG, where 14.8% and 13.3% of names map to multiple IDs; also in ChEBI ‘lecithin’ is the most ambiguous compound, linked to 413 IDs; other ambiguous names are, again, generic names such as ‘Diglyceride’, ‘Diacylglycerol’, ‘Triglyceride’ and ‘Triacylglycerol’ (see Figure 2A). Also in KEGG the most ambiguous names refer to generic compounds like ‘DS-18’ with 16 corresponding IDs. Further more this compound shares ID with ‘Chondroitin 4-sulfate’ which is a sulfated glycosaminoglycan while DS-18 generally refers to glycan, which further complicates metabolite characterization, as shown in Figure 2 B.

**Figure 2.**
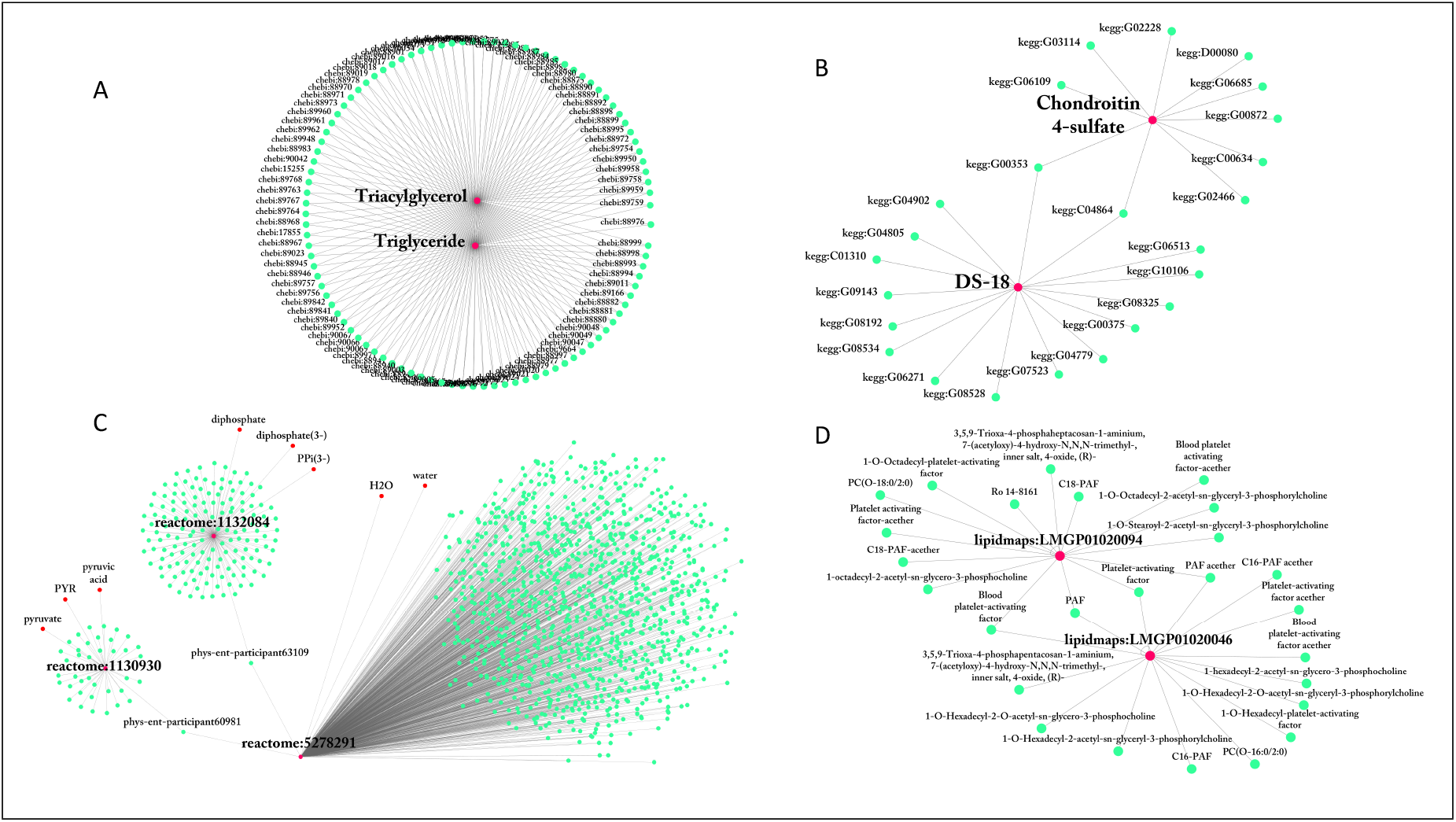
*Intra* database consistency. Edges indicate a link between a metabolite name and a database ID. Database name has been added to the ID (denoted as database names followed by ‘:’, *i.e* kegg:C00228). A-B: Examples of metabolite names associated with multiple IDs in ChEBI(A) and KEGG (B). C-D: Examples of metabolite IDs associated with multiple names in Reactome (C) and LIPID MAPS (D)

EnviPath and SLM databases have also relatively low consistency with 7% names being ambiguous. SLM is the largest database considered (> 1.2 × 10^6^ entries) and the most ambiguous name refers to ‘Triacylglycerol’. In enviPath the most ambiguous compound is ‘compound 0044249’, with SMILES representation CC1=CC=C(C=C1O)O that corresponds to 4-methyl-1,3-benzenediol. In this database, many metabolites are renamed with numbers, *i.e* ‘P06’,’M320I23’, or ‘compound 869’, which makes it cumbersome to the human user to identify them.

Other databases, namely, SABIO-RK, MetaCyc and LIPID MAPS are highly consistent, with SABIO-RK containing only 8 metabolites with ambiguous names.

#### 2.1.2 ID multiplicity and use of synonyms

In an effort to increase readability of entries in the database, often multiple names are linked to the same ID, i.e. IDs have a multiplicity larger than 1. Note that multiplicity is different from ambiguity as defined at the start of the Results section. Multiplicity increases human readability and is beneficial, as long as the alias, names and synonyms describe the same metabolite. Table 2 presents the average ID multiplicity for the eleven databases considered.

**Table 2.** ID multiplicity in each database: number of IDs in each database, average number of names per ID (average multiplicity), percentage and number of IDs that associate to more than one name, and highest number of names an ID links to.

| Database | #ID | Average multiplicity<br>± standard deviation | % of IDs with<br>multiplicity >1 | # of IDs with<br>multiplicity >1 | Highest multiplicity<br>in database |
| --- | --- | --- | --- | --- | --- |
| BiGG | 5174 | 1.0 ± 0.0 | 0.0 | 0 | 1 |
| ChEBI | 123835 | 4.3 ± 3.6 | 97.74 | 121034 | 57 |
| enviPath | 12306 | 1.0 ± 0.2 | 1.6 | 197 | 10 |
| HMDB | 43179 | 2.4 ± 0.5 | 99.71 | 43052 | 8 |
| KEGG | 40256 | 1.7 ± 1.2 | 38.93 | 15671 | 31 |
| LIPID MAPS | 40772 | 3.9 ± 1.0 | 100.0 | 40772 | 23 |
| MetaCyc | 17159 | 3.3 ± 2.0 | 99.75 | 17116 | 98 |
| Reactome | 5344 | 2.3 ± 16.6 | 47.46 | 2536 | 1106 |
| SABIO-RK | 7683 | 1.5 ± 1.2 | 24.17 | 1857 | 21 |
| SEED | 27693 | 1.7 ± 1.3 | 39.83 | 11031 | 28 |
| SLM | 505004 | 2.6 ± 0.6 | 99.87 | 504333 | 9 |

BiGG is the only exception to this rule. Every metabolite identifier is associated to one and only one metabolite name, but, as shown in Table 1, the contrary does not hold true. BiGG is the smallest database here considered (with only 5102 metabolite names and 5174 metabolite IDs), although it should be stressed that this database has been built by integrating reactions and metabolites appearing in several published and manually curated genome-scale metabolic networks.

All other databases have some extent of multiplicity: in ChEBI, HMDB, Metacyc, SLM and LIPID MAPS nearly 100% of IDs are linked to more than 1 name. The use of multiple names is intended to increase usability of the database. However, inconsistencies might arise when ambiguous names are linked to IDs with high multiplicity, as illustrated in Figure 2 C and D. This can result in errors and mismatches when identifying compounds. A most extreme case is Reactome identifier reactome:5278291 which is linked to 1106 difference names (see Figure 2C), among them ‘H2O’, ‘water’, ‘phys-ent-participant60981’ and ‘phys-ent-participant63109’. The latter two names are linked to identifiers pointing to ‘diphosphate’ and ‘pyruvate’, which means that within this database is possible to map ‘water’ to ‘pyruvate’. Other striking examples can be found in Table 3. When mapping with these compounds extra care needs to be taken.

**Table 3.**
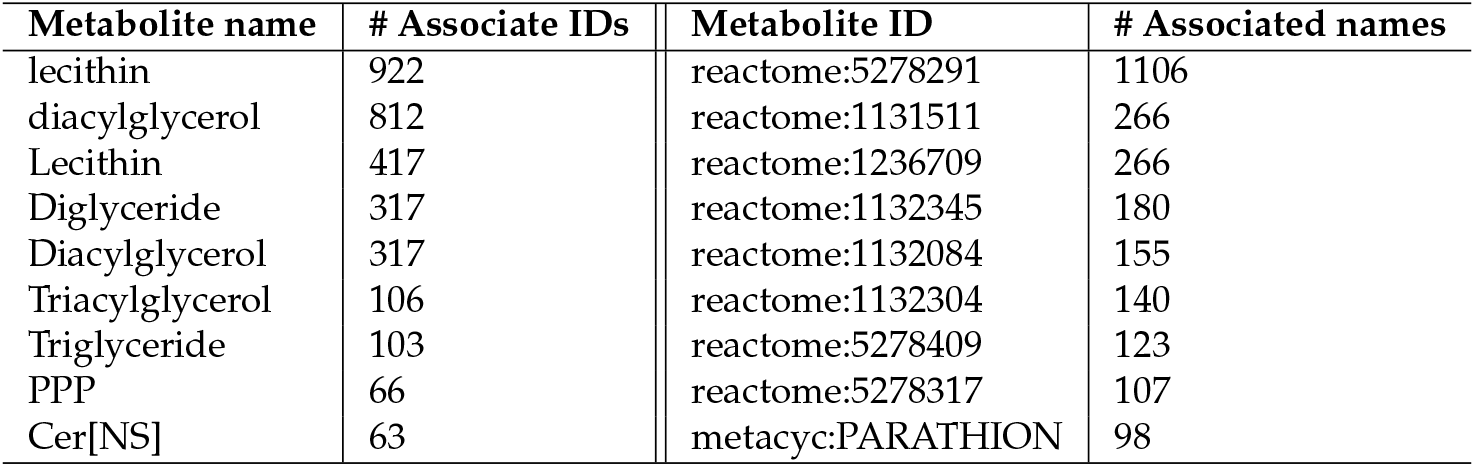
Example of compound names and IDs with high ambiguity and multiplicity.

| Metabolite name | # Associate IDs | Metabolite ID | # Associated names |
| --- | --- | --- | --- |
| lecithin | 922 | reactome:5278291 | 1106 |
| diacylglycerol | 812 | reactome:1131511 | 266 |
| Lecithin | 417 | reactome:1236709 | 266 |
| Diglyceride | 317 | reactome:1132345 | 180 |
| Diacylglycerol | 317 | reactome:1132084 | 155 |
| Triacylglycerol | 106 | reactome:1132304 | 140 |
| Triglyceride | 103 | reactome:5278409 | 123 |
| PPP | 66 | reactome:5278317 | 107 |
| Cer[NS] | 63 | metacyc:PARATHION | 98 |

#### 2.1.3. Database mapping to IDs from MNXRef

MNXRef is a common namespace derived from MetaNetX and has been developed to combine namespaces from multiple databases and provides links between compounds (and identifiers) from different databases, the overarching goal is to enable bringing together GEMs.

We found that, each of the IDs in the 11 databases link to a MNXRef ID, however, as shown in Table 4, one MNXRef ID can connect to several IDs within a database resulting in a multiplicity larger than 1. This happens, for instance, when MetaNetX associates one ID to several compound synonyms. This might be due to conscious modelling-specific decisions. For instance, it would make sense to combine citrate/citric acid identifiers in different databases to deal with protonation state differences. Thus linking several IDs to the same MNXRef ID addresses the multiplicity present in the database. However, this also generates errors if the ID links to ambiguous names. The most striking case is observed when mapping Reactome and MetaNetX: 5062 MetaNetX IDs are associated to Reactome IDs and 41.93% of them link to more than one Reactome ID.

**Table 4.** Number of IDs (#ID) in each database, number of MNXRef IDs (#MNXRef ID) linking to each database, multiplicity of MNXRef IDs when mapping to IDs in the corresponding database, and average and highest number of MNXRef ID per database ID

| Database | #ID | #MNXRef ID | average #ID per MNXRef ID<br>± standard deviation | % of IDs with<br>multiplicity >1 | # of IDs with<br>multiplicity >1 | Highest ID<br>multiplicity >1 |
| --- | --- | --- | --- | --- | --- | --- |
| BiGG | 5174 | 5062 | 1.0 ± 0.2 | 1.96 | 99 | 4 |
| ChEBI | 123835 | 96746 | 1.3 ± 1.0 | 11.93 | 11541 | 30 |
| enviPath | 12306 | 11087 | 1.1 ± 0.4 | 8.14 | 902 | 9 |
| HMDB | 43179 | 42354 | 1.0 ± 0.2 | 1.63 | 691 | 12 |
| KEGG | 40256 | 37722 | 1.1 ± 0.3 | 6.14 | 2316 | 12 |
| LIPID MAPS | 40772 | 40546 | 1.0 ± 0.1 | 0.51 | 207 | 6 |
| MetaCyc | 17159 | 16985 | 1.0 ± 0.1 | 0.9 | 153 | 5 |
| Reactome | 5344 | 2058 | 2.6 ± 3.9 | 41.93 | 863 | 34 |
| SABIO-RK | 7683 | 7512 | 1.0 ± 0.2 | 2.2 | 165 | 3 |
| SEED | 27693 | 26894 | 1.0 ± 0.2 | 2.79 | 749 | 4 |
| SLM | 505004 | 504881 | 1.0 ± 0.0 | 0.02 | 119 | 3 |

### 2.2. Namespace mapping between databases

To study namespace consistency between databases, we performed a pairwise mapping of the 11 databases. We performed the mapping using both the names in the corresponding database and MNXRef identifiers.

#### 2.2.1. Mapping between databases using metabolite names

Table 5 shows the results of pairwise database mapping using metabolite names. Here, we map IDs in the databases using associated names. The databases have different metabolite coverages, for instance SLM contains 1218750 names while BiGG only 5102, this is because some are specific for a certain class of compounds (like SLM for lipids) while others aim to be comprehensive and do not describe all compound classes in exhaustive details (like HMDB for lipds). The difference in coverage and multiplicity of names associated to IDs (previously presented in Table 1 and Table 2) can cause the mapping between two databases not to be symmetric as evident from Table 5.

**Table 5.** Number of IDs in one database (column) that map to IDs in the database in the corresponding row using database names as a bridge for mapping. Percentages indicate fraction of the initial database. Blue boxes indicate highest overall map

| Database | BiGG | ChEBI | enviPath | HMDB | KEGG | LIPID MAPS | MetaCyc | Reactome | SABIO-RK | SEED | SLM |
| --- | --- | --- | --- | --- | --- | --- | --- | --- | --- | --- | --- |
| BiGG | – | 5097 (4.1%) | 150 (1.2%) | 702 (1.6%) | 1489 (3.7%) | 158 (0.4%) | 210 (1.2%) | 361 (6.8%) | 839 (10.9%) | 1829 (6.6%) | 61 (0.0%) |
| ChEBI | 1303 (25.2%) | – | 816 (6.6%) | 9178 (21.3%) | 16013 (39.8%) | 4662 (11.4%) | 7209 (42.0%) | 2146 (40.2%) | 2552 (33.2%) | 15837 (57.2%) | 4336 (0.9%) |
| enviPath | 142 (2.7%) | 2284 (1.8%) | – | 304 (0.7%) | 1111 (2.8%) | 55 (0.1%) | 31 (0.2%) | 90 (1.7%) | 300 (3.9%) | 983 (3.5%) | 6 (0.0%) |
| HMDB | 643 (12.4%) | 15749 (12.7%) | 310 (2.5%) | – | 4745 (11.8%) | 4078 (10.0%) | 1693 (9.9%) | 877 (16.4%) | 1268 (16.5%) | 3868 (14.0%) | 14007 (2.8%) |
| KEGG | 1286 (24.9%) | 30098 (24.3%) | 1050 (8.5%) | 3922 (9.1%) | – | 1725 (4.2%) | 731 (4.3%) | 928 (17.4%) | 2604 (33.9%) | 16646 (60.1%) | 84 (0.0%) |
| LIPID MAPS | 149 (2.9%) | 7832 (6.3%) | 54 (0.4%) | 4200 (9.7%) | 1862 (4.6%) | – | 622 (3.6%) | 311 (5.8%) | 377 (4.9%) | 1893 (6.8%) | 13478 (2.7%) |
| MetaCyc | 212 (4.1%) | 20183 (16.3%) | 31 (0.3%) | 1967 (4.6%) | 851 (2.1%) | 648 (1.6%) | – | 1266 (23.7%) | 340 (4.4%) | 7703 (27.8%) | 326 (0.1%) |
| Reactome | 156 (3.0%) | 5833 (4.7%) | 41 (0.3%) | 620 (1.4%) | 588 (1.5%) | 254 (0.6%) | 717 (4.2%) | – | 368 (4.8%) | 542 (2.0%) | 146 (0.0%) |
| SABIO-RK | 864 (16.7%) | 10413 (8.4%) | 324 (2.6%) | 1456 (3.4%) | 3127 (7.8%) | 390 (1.0%) | 342 (2.0%) | 781 (14.6%) | – | 2692 (9.7%) | 55 (0.0%) |
| SEED | 1824 (35.3%) | 32212 (26.0%) | 1020 (8.3%) | 4971 (11.5%) | 18489 (45.9%) | 1915 (4.7%) | 7580 (44.2%) | 985 (18.4%) | 2641 (34.4%) | – | 233 (0.0%) |
| SLM | 55 (1.1%) | 4964 (4.0%) | 4 (0.0%) | 12354 (28.6%) | 94 (0.2%) | 10634 (26.1%) | 289 (1.7%) | 225 (4.2%) | 44 (0.6%) | 211 (0.8%) | – |

In all comparisons, the fraction of compounds sharing the same name is rather limited. Overall, except for mapping from SEED to KEGG and ChEBI with 60.1 % and 57.2% overlap, respectively, all databases have less than 50% of compound names in common. The namespace of ChEBI has the largest overlap with other namespaces: around 40% towards MetaCyc, Reactome, and KEGG can be mapped to ChEBI. The namespaces of SLM, enviPath, and LIPID MAPS have the smallest overlap with other namespaces, which is most likely because these are very specific databases. The low ratios in Table 5, indicate that mapping using string algorithms is not effective since trivial differences in the names (such as the use of underscore and hyphen) can results in mismatches.

Ambiguous naming, *i.e*. one name associated to more than one ID, can result in mapping inconsistencies where one ID in the first database, gets mapped to multiple IDs in the second database. The fraction of non-univocal mappings is indicated in Table 6. Hence, although 40.2% of the Reactome IDs can be mapped to ChEBI (see Table 5), 81.2% of the successfully mapped Reactome IDs are ambiguously mapped to multiple ChEBI IDs.

**Table 6.** Percentage of IDs in the database in the column that get mapped to more than one ID in the database in the corresponding row using database names as a bridge. Blue boxes are used to highlight highest numbers.

| Database | BiGG | ChEBI | enviPath | HMDB | KEGG | LIPID MAPS | MetaCyc | Reactome | SABIO-RK | SEED | SLM |
| --- | --- | --- | --- | --- | --- | --- | --- | --- | --- | --- | --- |
| BiGG | – | 2.9 | 1.3 | 3 | 3.6 | 3.2 | 1.4 | 0.6 | 2.9 | 2.7 | 1.6 |
| ChEBI | 76.3 | – | 67 | 38.1 | 38.3 | 34.3 | 58.7 | 81.3 | 78.7 | 37.3 | 26.9 |
| enviPath | 6.3 | 6.5 | – | 8.2 | 6.1 | 0 | 0 | 12.2 | 7.7 | 4.6 | 0 |
| HMDB | 10.7 | 11.5 | 6.8 | – | 7.3 | 4.3 | 13.2 | 22.8 | 12.8 | 7.4 | 0.7 |
| KEGG | 17 | 15.2 | 11.1 | 28.5 | – | 10.2 | 18.5 | 34.5 | 19.6 | 12.4 | 33.3 |
| LIPID MAPS | 8.7 | 9.8 | 1.9 | 1.8 | 3.2 | – | 4.2 | 13.2 | 4.5 | 3.2 | 0.8 |
| MetaCyc | 0.5 | 3.9 | 0 | 2.5 | 3.9 | 2 | – | 6 | 4.1 | 1.5 | 0.6 |
| Reactome | 42.3 | 41.4 | 51.2 | 49 | 51.4 | 24.4 | 38.9 | – | 49.5 | 43.2 | 47.9 |
| SABIO-RK | 0 | 4.5 | 0 | 0 | 3.8 | 1 | 3.8 | 2.2 | – | 3.3 | 1.8 |
| SEED | 3 | 6 | 0.9 | 2 | 2.4 | 2.2 | 3.1 | 8.9 | 5.3 | – | 1.7 |
| SLM | 7.3 | 37.2 | 25 | 12.3 | 18.1 | 22.3 | 10.4 | 24.4 | 20.5 | 9.5 | – |

In some cases, more than 50% of the mappings are non-unique. The highest fractions of non unique ID mapping occurs when mapping to ChEBI, although when mapping from ChEBI to the other databases, this fraction reduces significantly. When considering Reactome, both mappings to and from this database lead to relatively high number of non-univocal assignments. SLM and SABIO-RK have a significant low ambiguity when mapping from other databases, although as shown in Table 5, only a small fraction in these databases can be mapped from other databases.

#### 2.2.2. Mapping between databases using MNXRef ID

Another approach to map IDs from different databases is to use MetaNetX/MNXRef as a bridge. Table 7 shows the fraction of IDs in each database pair that can be mapped through MetaNetX/MNXRef. Again the differences in coverage between the databases cause this table to be non-symmetric.

**Table 7.**
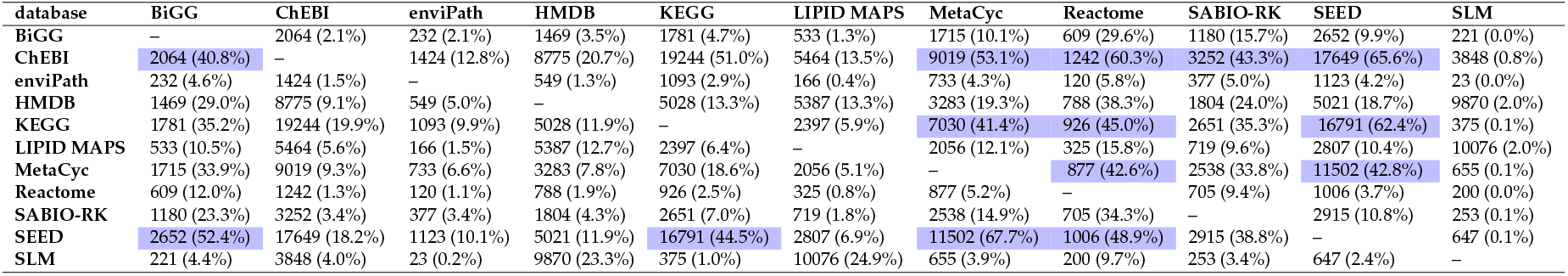
Number of IDs in one database (column) that map to IDs in the database in the corresponding row using MetaNetX as a bridge. Percentages indicate fraction of IDs in the initial database. Blue boxes are used to highlight highest numbers.

It is clear that this approach results in more identified mappings than the previous approach that used names. Nevertheless, the overall map is also not high. None of tested databases maps higher than 70 % either to or from other databases. The highest match is 67.7 % when mapping MetaCyc to SEED. SEED can be mapped fairly well from BiGG, Reactome and KEGG with more than 40% match. Note that these are all databases specialized in reactions and metabolic pathways. There is almost no overlap between SEED and SLM, the latter specialized in lipids. Databases with overall good match are ChEBI, KEGG and MetaCyc. Among them, ChEBI has the highest overlap with other databases. Almost 50 % of IDs in SEED, Reactome, MetaCyc, KEGG, SABIO-RK and BiGG can be mapped to ChEBI. However, there is not so much overlap when mapping enviPath (12.8%), LIPID MAPS (13.5%) and especially SLM (0.8%) to ChEBI. The remaining databases have a significant low overlap percentage when mapping via IDs. Especially SLM, there is just a minor part of the database that can be mapped to other databases.

This approach also results in instances of one ID from the first database associated to more than one ID in the target database, an example is provided in Figure 3 and Table 8 summarizes the identified cases.

**Figure 3.**
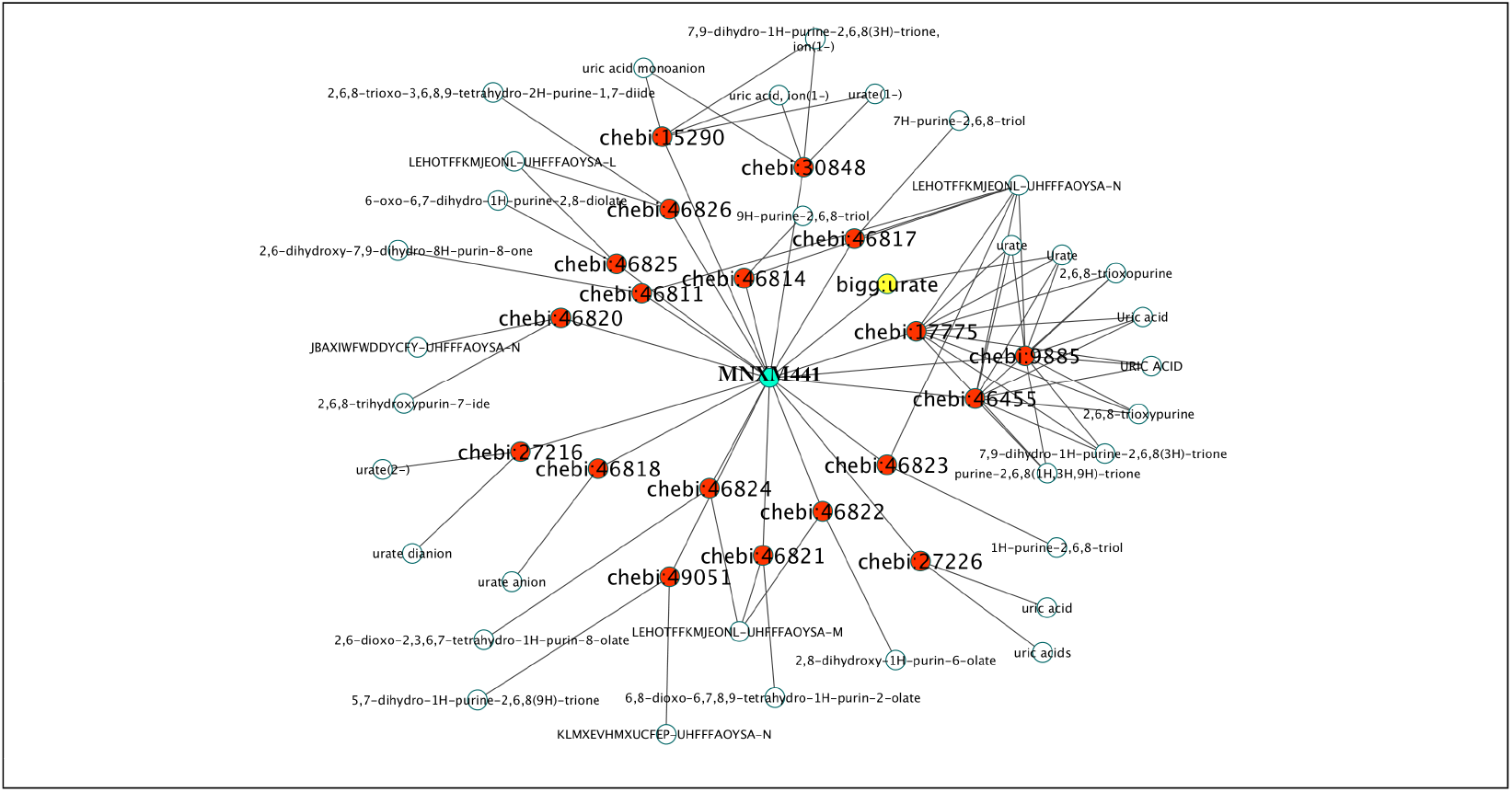
Visualization of the *inter* database inconsistency. An ID from BiGG (in yellow) can link to many other IDs in CheBI (red) when using MetaNetX ID (green) for the mapping.

**Table 8.** Percentage of IDs in the database in the column that get mapped to more than one ID in the database in the corresponding row using database MetaNetX as a bridge. Blue boxes are used to highlight highest numbers

| Database | BiGG | ChEBI | enviPath | HMDB | KEGG | LIPID MAPS | MetaCyc | Reactome | SABIO-RK | SEED | SLM |
| --- | --- | --- | --- | --- | --- | --- | --- | --- | --- | --- | --- |
| BiGG | – | 3.9 | 5.2 | 3.5 | 3.9 | 3.2 | 4 | 3.8 | 4.5 | 3.2 | 2.7 |
| ChEBI | 83.1 | – | 56.2 | 39.7 | 36.4 | 37.8 | 64.7 | 76.8 | 72.2 | 39.4 | 27.8 |
| enviPath | 9.9 | 10.6 | – | 12 | 8.1 | 8.4 | 7.6 | 14.2 | 11.1 | 8.1 | 8.7 |
| HMDB | 19.1 | 6.8 | 12.6 | – | 9.3 | 5.1 | 12.7 | 26.4 | 17.2 | 9.7 | 1.6 |
| KEGG | 15 | 10 | 11 | 22.1 | – | 8.4 | 9.6 | 19.7 | 17.5 | 11.2 | 14.7 |
| LIPID MAPS | 10.5 | 2.8 | 6 | 2.5 | 4.5 | – | 4.6 | 14.5 | 7 | 4.2 | 0.5 |
| MetaCyc | 3.6 | 1.4 | 3.4 | 2.1 | 1.7 | 0.9 | – | 4.3 | 2.8 | 1.1 | 0.5 |
| Reactome | 42.7 | 33.3 | 45 | 32.4 | 37.1 | 28.9 | 36.7 | – | 41.7 | 37.2 | 35 |
| SABIO-RK | 8.1 | 4.7 | 5.6 | 6.1 | 5.3 | 3.8 | 5.6 | 9.2 | – | 5.1 | 4.7 |
| SEED | 8.4 | 3.5 | 4.2 | 4.6 | 3.7 | 3.6 | 5.7 | 12.1 | 9.5 | – | 5.1 |
| SLM | 5 | 1.1 | 0 | 0.2 | 5.6 | 0.4 | 2.7 | 6 | 5.1 | 3.2 | – |

Name ambiguity and non-unique ID mapping between databases can lead to inconsistencies (different metabolites being considered to be equivalent) and included as such in the metabolic model.

Table 9 lists some illustrative examples. These examples show that automatic mapping (manual mapping is impossible for large scale models) of compounds between or within databases can lead to introduction of unrealistic reactions that can potentially reduce the accuracy of the predictions of the model.

**Table 9.**
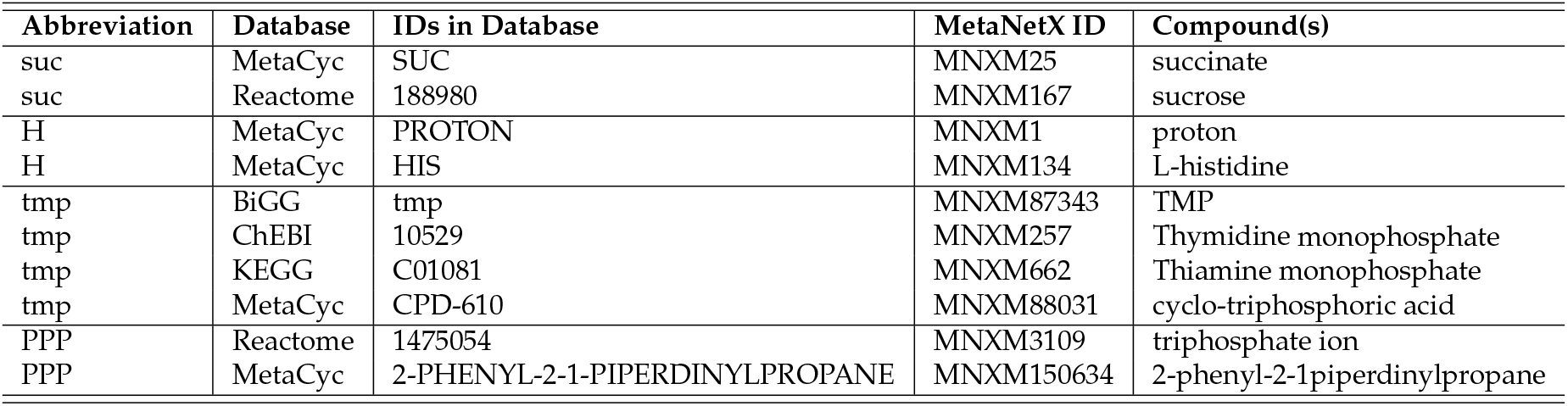
Examples of mapping inconsistencies

## 3. Discussion

GEMs aim to be comprehensive representations of the metabolism of one organism. They are often built based on more than one database. As explained, the initial step of model constructions is typically automated model drafting. Tool selection will determine which name space the model is associated to. For instance, modelSEED uses SEED as a reference reaction database while Pathway Tools uses MetaCyc. In the next step in the model building process - manual curation - gap filling is possibly the most important task. Tools for gap-filling often systematically explore the GEM to identify possible gaps [33], Other methods rely on additional experimental data such as measured metabolites to identify the gaps [34]. In this step, researchers may use different sources and databases to identify reactions and associated metabolites. Errors might arise due to inconsistencies in this mapping A second application of GEMs is the integration and contextualization of ‘omics’ data such as transcriptomics, proteomics, metabolomics and/or fluxomics data. These applications often require a mapping of metabolite identifiers to match the namespace of the model and that of the database that has been used in the data generation process. Both applications may imply potential problem(s) caused by ambiguous names or identifiers.

Among the eleven explored databases, KEGG, BiGG, ChEBI, MetaCyc, HMDB and SEED are the most commonly used in metabolic modeling. We calculated the ambiguity of names and the multiplicity of non-systematic identifiers within and between eleven databases. Within the same database, the percentage of identifiers with multiplicity larger than one varies from 0 % to 100 %, whereas the ambiguity of names ranges from 0.07 % to 29.4 %. When mapping between databases, these ambiguities and multiplicities lead to larger inconsistencies, and this agree with previous observations regarding small molecules databases [31,35]. The inconsistencies when mapping using metabolite names range from 0 % to 81.2 %. Similar results are obtained while mapping via MNXRef ID, between databases, as the number of inconsistencies varies from 0 % to 83 %, however on average better results are obtained. Mapping with the databases with the highest number of ambiguous names also results in higher number of inconsistencies than when mapping between other databases. Among the eleven tested databases, Reactome, HMDB, ChEBI, and KEGG are those that show the highest *intra*- and *inter*-database ambiguity.

Most of the ambiguous names are associated to general compounds such as triacylglycerol, glycan or protein. These names and IDs represent classes of compounds rather than metabolites with defined structures and are included in metabolic models as they have a clear biological interpretation. However, care should be taken when introducing them in databases and these names should not be included in the list of synonyms for specific compounds, as mentioned in [35]. Using abbreviations to refer to compounds is also highly ambiguous as the same abbreviations can represent different compounds in the same or in different databases.

Our findings show that compound names or IDs cannot be clearly mapped automatically. Even if we use non-ambiguous identifiers, many mappings are still inconsistent because they can link to ambiguous names. MetaNetX solved some of the issues as shown in Table 9. However, not all compounds in the eleven tested databases can be mapped with MNXRef. Mapping from MetaCyc to SEED, SEED to ChEBI, and SEED to KEGG using MNXRef give the highest number of matches, but still only around 60 % of compounds matched. Other databases show much lower coverage.

In order to use metaNetX/MNXRef ID to map compounds in a GEM, the namespace of the model needs to be related to at least one of the eleven databases considered. However, many models uses custom made naming conventions [36]. For these models, mapping through name is the only option.

Ambiguous namespaces also hamper the (re)use of models from different research labs or organisms. Due to a low level of interoperability, in practice it is impossible to directly compare models, as metabolites can hardly be cross-mapped, which in turn makes it impossible to compare reactions in both models, see examples in Table 9. Nevertheless, comparing models is important and necessary: it helps to reduce the time to build models for closely related species; to combine efforts from different research groups that study the same organism; and to study the metabolic differences between different organisms. In addition, microbial communities are notoriously difficult to characterize. While transcriptomics and proteomics measurements can be associated (to a great extent) to the originating microorganism, it is not possible to do this for metabolites. Therefore, there is a need for models that can help combine both types of measurements. As a result, there are on-going efforts to define modelling frameworks, based on combining GEMs of individual organisms, to characterize the behaviour of the community [21,22,37,38]. Enabling unambiguous mapping will be required to take full advantage of these on-going developments.

Below we have enumerated a number of recommendations that may increase the level of interoperability of GEMs, facilitating unambiguous mapping

- Limit the use of aliases, *i.e*. compound classes or abbreviations, as synonyms in databases. These aliases increase human readability, but should be clearly distinguished from names and synonyms in the databases and should not be used for mapping.
- In the context of metabolic modelling it is frequent and desirable to use compound classes to identify generic compounds [28]. Compounds like ‘biomass’ or ‘lipid’ are often used in GEMs; this does not affect the use of the model, except when predicting or simulating the production (of a specific component) of generic compounds, *i.e*. when ‘lipids’ are the main focus of the model. In fact, it is often better to use generic compounds whenever a specific compound is not needed, as they can be universal. For instance, ‘biomass’ has been used as a standard among the modelling community as an artificial compound that represents the growth objective of the cell [17,39]. Another reason is that often the precise identity of the compounds is not needed and there is a lack of experimental data for their characterization. Therefore, when using generic compounds, it is desirable to add extensive annotation to the model to clearly state which compounds they represent and for which purpose they are used in the model. These generic compounds are among the most ambiguous entities in the eleven analyzed databases and we therefore advise to exclude them from any automatic mapping process.
- Avoid using highly ambiguous names as the sole description of the compound in the model. When referring to these compounds, clear annotation needs to be included to prevent mismatches and inconsistencies.
- In addition to human-readable identifiers and database-dependent identifiers, include database-independent identifiers, *i.e*. InChI [15,16] whenever possible for compounds with defined structures. Using InChI can help to fully automate the mapping [28]. However, take into account that mismatches and errors can also happen because identifiers can also link to incorrect InChI as shown in [29,35,40].
- Model mapping only based on metabolite information can imply certain mismatch due to differences in namespaces, even if systematic identifiers were used. Hence, different mapping strategies, *i.e*. mapping through encoding genes and network topology [19], should be used to complement name or identifier based mapping.
- GEMs also need to have a unique standard annotation so that they generate the same output even when different tools are used for the simulation. Neal *et al*. [41] suggest that semantic annotation can help to store and combine models, but these models need to stick to a unique standard annotation format.

Simply deciding a standard database/identifier/annotation to represent metabolites in models will also not help to improve the situation, as they will limit the available model construction tools. Nevertheless, while increasing the level of interoperability none of the presented approaches above can by itself ensure automated mapping without errors. Different approaches need to be combined when translating between namespaces. Manual curation is still required, at least for compounds with highly ambiguous names.

Finally, our analysis has some limitations. We have only studied non-systematic identifier and names. We did not use structure data such as MOL files, hence we cannot identify all incorrect matches. We have not included such information in our analysis because it is not often found in metabolic models. In addition, MetaNetX data that was downloaded at the moment of conducting this study contained data from the originating databases that was produced in 2017 and some in 2016 (see section 4.3 for more detail). As databases change over time, a similar analysis with the most recent database updates might lead to different results. *Stat Roma pristina nomine, nomina nuda tenemus*.

## 4. Material and Methods

### 4.1. Data collection and preprocessing

Data about compound identifiers and synonyms were downloaded from MetaNetX[25]. MetaNetX is a repository of GEMs and biochemical pathways. It contains entries from some of the most relevant databases that have been used in GEMs construction and simulation such as KEGG, BiGG, MetaCyc and SEED [42]. The platform (http://www.metanetx.org/) allows access to these databases as well as provides tools to map/translate them. In this study, The chem_xref.tsv file was downloaded from the MetaNetX website on 31^*st*^ October, 2018. In the following, we provide a brief description of the content of these databases.

Biochemical, Genetic and Genomic models (BiGG) [43] is a knowledge database of genome scale metabolic models (GEMs). Currently, it contains 85 high-quality, manually curated GEMs, 24311 reactions, and 7339 metabolites (data retrieved on 30^*th*^, Nov, 2018 from http://bigg.ucsd.edu/). In BiGG, the metabolite is identified as the abbreviation of its name. For example, ‘10fthf’ for 10-Formyltetrahydrofolate. MetaNetX obtained data from BiGG on 2017/04/11.

Model SEED [44] is a platform to construct GEMs that uses its own database for metabolites and reactions. This database combines information from KEGG and existing metabolic models in a non-redundant set of reactions. In this database, metabolite identifiers start with “cpd” and followed by a 5 digits number. For example, D-Glucose-1-Phosphate is cpd00089. The database can be downloaded from https://github.com/ModelSEED/ModelSEEDDatabase/tree/master/Biochemistry.MetaNetX obtained data from SEED on 2017/04/13.

ChEBI [45]. (http://www.ebi.ac.uk/chebi/aboutChebiForward.do) is a database of Chemical Entities of Biological Interest [45] and is a repository for small chemical compounds. In ChEBI, metabolites are named by 5 digit numbers. For example, Alpha-D-glucose-1-phosphate(2-) is 58601. File can be downloaded from ftp://ftp.ebi.ac.uk/pub/databases/chebi/Flat_file_tab_delimited/. ChEBI data in MetaNetX are from the release version 150. enviPath [46]. (https://envipath.org/). Is a database to store and predict the microbial biotransformation of organic environmental contaminants. Data in MetaNetX were downloaded on 2017/04/12.

HMDB [47], (http://www.hmdb.ca), is a comprehensive and curated collection of human metabolite and human metabolism data. Data in MetaNetX was obtained on 2017/04/12.

KEGG [13] (http://www.KEGG.jp). The Kyoto Encyclopedia of Genes and Genomes is a resource that provides information about pathways and reactions in organisms. In KEGG, metabolites started with a letter ‘C’ (compound) and followed by 5 digit numbers. For example, D-Glucose-1-Phosphate is identified as C00103. Data in MetaNetX were obtained on 2017/04/12.

LIPID MAPS [48]. (http://www.lipidmaps.org). Is a database that contains structures and annotations of biologically relevant lipids. Data in MetaNetX were obtained on 2017/04/13.

MetaCyc [12]. (http://metacyc.org). Is a curated database of metabolic pathways. All data in MetaCyc are experimentally validated. The metabolite is identified by its full name. For example, D-glucose-1-Phosphate is D-glucose-1-phosphate. The database can be downloaded here http://bioinformatics.ai.sri.com/ptools/flatfile-format.html. Data in MetaNetX were obtained on 2017/04/13.

Reactome [49]. (http://www.reactome.org). Is a curated and peer-reviewed database of human biological processes. Data in MetaNetX were obtained on 2017/04/13.

SABIO-RK [50], (http://sabiork.h-its.org/), Is a database containing comprehensive information about biochemical reactions and their kinetic properties. Data in MetaNetX were obtained on 2016/05/27.

SwissLipids (SLM) [51] (http://www.swisslipids.org/) contains curated data about lipid structures and metabolism. Data in MetaNetX were obtained on 2017/04/13.

The original data file was modified prior to analyzing. The modification includes the removal of the description part, of IDs starting by bigg:M as they are not real compound ID in BiGG, and the removal of ‘biomass’ compounds. Data from MetaNetX were organized in four columns in this order: compound ID in original database with database indicator in front, for example bigg:10fthf, corresponding compound IDs in MetaNetX, evidence and description (name).

### 4.2. Intra-database analysis

For intra-database consistency analysis, the first, the second and the last column of the MetaNetX data file were used for mapping. Name ambiguity was calculated as the number of ID each name links to. Similarly, the name multiplicity of each ID was calculated as the number of names it refers to.

### 4.3. Inter-database analysis

We mapped compound IDs between databases. A direct map between IDs in the database is not possible. The tested databases use different system for compound identifiers. For instance, in KEGG, the compound ID is a capital ‘C’ following by a 5 digit numbers, *i.e*. ‘C00002’ for ATP. In contrast, in BiGG, the compound is identified as abbreviation of its name, for example, ‘atp’ for ATP. Therefore, to map from one ID in database 1 to other ID in database 2, we used either the associated compound name or the associated MNXRef ID. That is also what MNXRef is meant for, as a link between databases.

Mappings via name were done by link from name to name in one database to the other. However, string algorithms are intolerant to even small changes such as the use of underscores or brackets. We first identified all compound IDs from one database. From this list, we counted the number of IDs that link to a name that is shared with the other database.

## Acknowledgments

This work has received funding from the Research Council of Norway, No. 248792 (DigiSal) and from the European Union FP7 and H2020 under grant agreements No. 305340 (INFECT), No. 635536 (EmPowerPutida) and No. 730976 (IBISBA 1.0)

## Author Contributions

NP, RvH, JvD and MSD designed and conceived the experiments. NP performed the data retrieval and analysis. NP, RvH, PS, ES and MSD performed the data interpretation. NP drafted the manuscript. ES, RvH and MSD edited the manuscript. All authors read and approved the manuscript.

## Conflicts of Interest

The authors declare no conflict of interest. The founding sponsors had no role in the design of the study; in the collection, analyses, or interpretation of data; in the writing of the manuscript, and in the decision to publish the results.

